# Plug-and-Play Myoelectric Control via a Self-Calibrating Random Forest Common Model

**DOI:** 10.1101/2024.11.07.622455

**Authors:** Xinyu Jiang, Chenfei Ma, Kianoush Nazarpour

## Abstract

**Objective:** Electromyographic (EMG) signals show large variabilities over time due to factors such as electrode shifting, user behaviour variations, etc., substantially degrading the performance of myoelectric control models in long-term use. Previously one-time model calibration was usually required each time before usage. However, the EMG characteristics could change even within a short period of time. Our objective is to develop a self-calibrating model, with an automatic and unsupervised self-calibration mechanism.

**Approach:** We developed a computationally efficient random forest (RF) common model, which can (1) be pre-trained and easily adapt to a new user via one-shot calibration, and (2) keep calibrating itself once in a while by boosting the RF with new decision trees trained on pseudo-labels of testing samples in a data buffer.

**Main results:** Our model has been validated in both offline and real-time, both open and closed-loop, both intra-day and long-term (up to 5 weeks) experiments. We tested this approach with data from 66 able-bodied participants. We also explored the effects of bidirectional user-model co-adaption in closed-loop experiments. We found that the self-calibrating model could gradually improve its performance in long-term use. With visual feedback, users will also adapt to the dynamic model meanwhile learn to perform hand gestures with significantly lower EMG amplitudes (less muscle effort).

**Significance:** Our random forest-approach provides a new alternative built on simple decision tree for myoelectric control, which is explainable, computationally efficient, and requires minimal data for model calibration. Source codes are avaiable at: https://github.com/MoveR-Digital-Health-and-Care-Hub/self-calibrating-rf

## 1. Introduction

Myoelectric control systems translate electromyographic (EMG) signals to control commands of human-machine interfaces [1, 2, 3, 4], enabling users to interact with diverse devices, e.g. prosthesis or exoskeleton in clinical applications [5, 6, 7, 8, 9], or, virtual keyboard and menu navigation in mixed reality applications [10, 11]. The variability of EMG patterns over time (both intra- and inter-day) is a key problem limiting the practical applications of myoelectric control systems. Various complicated factors, such as noises, the behaviour variation of users, muscle fatigue, electrode shifting, limb positions and other unknown physiological factors jointly result in the EMG variability [12, 13, 14]. In real-world applications, user motor learning (even in open-loop myoelectric control [15]) can also gradually change EMG patterns over time. The EMG variability substantially degrading the performance of myoelectric control models in long-term use.

Previous studies aim to address the above problem by domain adaptation and transfer learning. These techniques can be further divided into three categories, namely supervised, semi-supervised, and unsupervised learning. As for domain adaptation, Vidovic et al. developed a covariate shift adaptation approach to align the statistical metrics of both training and testing data [16]. Guo et al. proposed an unsupervised domain adaptation approach based on locality preserving and maximum margin (LPMM) criterion, to automatically adapt the myoelectric control model to a shifted data distribution in different data collection sessions [17]. Shi et al. developed a multi-task dual-stream supervised domain adaptation (MDSDA) approach to improve the long-term robustness of myoelectric control models [18]. Transfer learning-based approaches could also effectively improve the performance of myoelectric control models by pre-training and fine-tuning a neural network in a supervised way [19]. Previous studies have also applied semi-supervised [20] and unsupervised learning [21, 22] to achieve the self-training or self-calibration of a deep learning model. A day-to-day model re-calibration algorithm incorporating semi-supervised learning and maximum independence domain adaptation (MIDA) has also been employed to improve the model generalisability [23]. In recent studies, the domain-adversarial neural network was employed to better generalise a model via an adversarial training manner [24, 25]. These previous efforts largely improve the long-term robustness of myoelectric control models.

However, previous studies mainly mitigated the above problems by performing one-time model calibration each time the model was applied. Furthermore, most previous studies performed offline validations by pooling together all testing samples in each session to evaluate the average performance. In real world scenarios, the EMG patterns may change slowly over time even in the same experimental session, which is a highly dynamic process. The continuously changing EMG characteristics remain a challenge for real world myoelectric control applications, but at the same time, also provide sufficient intermediary data serving as a bridge for our model to gradually track the variations and better generalise to latest characteristics of EMG. Additionally, although the above transfer learning-based techniques can compensate for the performance degradation due to the mismatch between training and testing data, sufficient training data with ground truth labels is still necessary to build the initial model. With the above-unsolved challenges, we asked: “Is it possible to develop a plug-and-play myoelectric control model efficiently (e.g. via one-shot learning), and enable the model to keep calibrating itself once in a while to track the slowly changing EMG characteristics?”

In this work, we develop a self-calibrating random forest (RF) model, which (1) can be pre-trained on other users and efficiently adapt to a new target user through one-shot calibration, built upon the method proposed in our recent work [26] and (2) self-calibrate *once in a while* based on pseudo-labels of testing samples saved in a data buffer. The pseudolabels used in the self-calibration process were assigned jointly by unsupervised manifold learning and clustering. Importantly, the whole framework in our work is built on decision trees, which are explainable, parallelisable and computationally efficient without too many floating point computations. The effectiveness of the overall framework was validated in both offline and real-time experiments, in both open-loop and closed-loop settings, and in intra-day, inter-day (2 consecutive days) and long-term (up to 5 weeks) applications. Instead of performance degradation over time, the self-calibrating RF model gradually improves its performance and keeps it at a stable level in long-term applications. For the dynamic self-calibrating model, we also explore the effects of bidirectional user-model co-adaption in closed-loop experiments. With visual feedback, users can also adapt to the dynamic self-calibrating model and learn to perform the target hand gestures with significantly lower EMG amplitudes (help users save efforts).

## 2. Methods

### 2.1. Experiments

#### 2.1.1. Ethical Approval

All participants joined our study voluntarily and gave written informed consent. Our study was approved by the ethics committee at the University of Edinburgh (reference number: 2019/89177), in adherence to the principles outlined in the Declaration of Helsinki. We conducted three experiments involving a total of 66 able-bodied subjects. Experiments 1 and 2 were offline while experiment 3 was run in real-time.

#### 2.1.2. Experiment 1

In experiment 1, we recruited 20 participants (12 males, 8 females, aged 22–43 years). Eight Delsys Trigno sensors were equally spaced across the circumference of the forearm as shown in Figure 1), which were also the common eight electrodes used to analyze data in all following experiments. Note that in experiment 3, sixteen electrodes were placed to record signals with a double electrode density (to support more diverse research purposes in the future), but all model calibration/testing steps were still performed using the same eight electrodes as shown in Figure 1. Sensors were configured with 10–500 Hz passband and a 2000 Hz sampling rate. Each participant performed 6 hand gestures listed in Figure 1. These hand gestures are the grip patterns that are quite useful for prostheses. A number of previous studies [27, 28] selected the same or a similar set of hand gestures. For each hand gesture, 10 repetitions were performed in 10 successive trials with a 6s duration for each trial. A 5s inter-trial resting period was provided. For each trial, data recorded in the first 2s reaction and transition period were removed.

**Figure 1:**
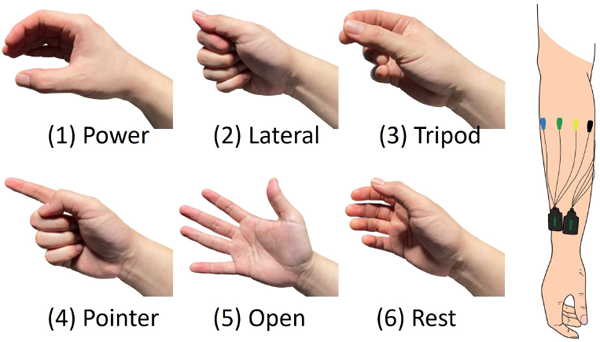
Hand gestures and electrode setup in our experiment.

#### 2.1.3. Experiment 2

In experiment 2, we recruited 18 *new* participants (11 males, 7 females, aged 22–28 years). Data were collected on two successive days. The experiment on day 1 comprised two sections, the calibration section and the testing section. In the calibration section, each participant performed only one repetition (in a 2s trial) per hand gesture. In each trial, participants would react to the experiment instruction program and shape their hands in one second, and hold the gesture in the next second. Only EMG signals recorded during the latter 1s period were retained (the same for all our following experiments) and used in our analyses. The testing section consisted of 5 testing blocks. In each testing block, participants performed 5 repetitions per hand gesture (a total of 30 repetitions), with a pseudo-randomised order. Participants were provided with 2s inter-trial and 5 minutes inter-block resting periods, respectively. On day 2, all participants started the testing section (the same as day 1) directly without the calibration section. Datasets collected in experiments 1 and 2 were used in offline analyses.

#### 2.1.4. Experiment 3

Experiment 3 was a real-time myoelectric control experiment. We recruited 28 *new* participants (aged 21–42 years, 13 males, 15 females). Experiment 3 was also conducted on two successive days. On day 1, the experiment consisted of a calibration section and a testing section, similar to experiment 2, except that, in each of the 5 testing blocks, data were unbalanced across different hand gestures. Specifically, each testing block did not necessarily involve exactly 5 repetitions per gesture. In this way, we can simulate a more realistic application scenario where data in a short period of time may be unbalanced, which is expected to shift the overall data distribution. We still keep data in 5 successive blocks on each day balanced across different hand gestures, so that we can reliably evaluate the model performance.

On day 1, models were calibrated immediately after the calibration section, and run in real-time in the testing section. We run two models in the testing section, one with and the other without self-calibration (details will be described later). The 5 testing blocks on day 1 were conducted in an open-loop mode, that is, participants did not see the output of both models. Therefore, any differences between the two models are completely due to their decoding capacity.

On day 2, the testing section directly started without a calibration section. The testing section on day 2 consisted of two parts, 5 testing blocks for each part. In part 1, the experiment is exactly the same as the testing section on day 1. After Part 1, all 28 participants were divided into two groups (Group A and Group B). Participants in both groups are with a similar average accuracy in part 1. After grouping each participant, the part 2 started. The experiment in part 2 for Group A was exactly the same as part 1 (open-loop mode), while the experiment for Group B was closed-loop with visual feedback. Specifically, participants in Group B were provided with the real-time outcome of the self-calibrating model together with the overall accuracy score of the model in each trial (shown at the end of a trial). The real-time outcome of the self-calibrating model was presented in the form of pictures of the decoded hand gestures, the same as the ones shown in Figure 1.

We randomly selected 2 participants in Group B to take our long-term experiment. The 2 participants came back each week (up to 5 weeks) to repeat our closed-loop experiment (exactly the same as part 2 on day 2).

### 2.2. Feature Extraction

In all experiments, features were extracted via a sliding window (window length: 200 ms; sliding step: 100 ms). In each window and each EMG channel, ten features were extracted to represent EMG signals from three complementary aspects. First, the mean absolute value (MAV), root mean square (RMS) waveform length (WL), slope sign changes (SSC), and zero crossings (ZC) were extracted as energy descriptors [29]. Second, the skewness of the EMG signals was calculated as a distribution descriptor [30, 31]. Third, the mean frequency (MNF), median frequency (MDF), peak frequency (PKF), and variance of central frequency (VCF) were used as spectrum descriptors [32]. The complementary role of the three types of features has been demonstrated in a previous study [26].

### 2.3. Pre-Training and Fine-Tuning a RF Model

#### 2.3.1. Pre-Training a RF Model

The RF model for each target user was pre-trained on all data recorded from other users. The pre-trained RF comprised 200 decision trees. Each feature from each subject was normalised separately. Considering the pre-training dataset is relatively large, 7% samples (suggested by [26]) were randomly drawn via bootstrap to train each decision tree, to avoid extremely complex decision rules and encourage diversity among decision trees.

#### 2.3.2. Fine-Tuning a RF Model

All pre-trained decision trees were pruned using the calibration data (one repetition per hand gesture, collected in the calibration section) from the new target user, denoted as *𝒮* _*target*_. If a node was pruned, all its children nodes (and children of children) were removed from the tree, and the pruned node would be a new leaf node. We applied a bottom-up pruning strategy. We first operated pruning on the parents nodes of the leaf nodes with the longest distance (*d*_*max*_) away from the root node. The calibration data with ground truth labels from the new user were used to validate the effect of each pruning operation. The pruning operation was performed only if the validation accuracy would not decrease without the pruned nodes. The pseudo-codes of decision tree pruning are presented in Algorithm 1. Details can be found in [26]. After inspecting the most distant lead nodes, we then inspected their parent nodes (moving to the top end until the root node).

##### Algorithm 1 Bottom-Up Tree Pruning Strategy

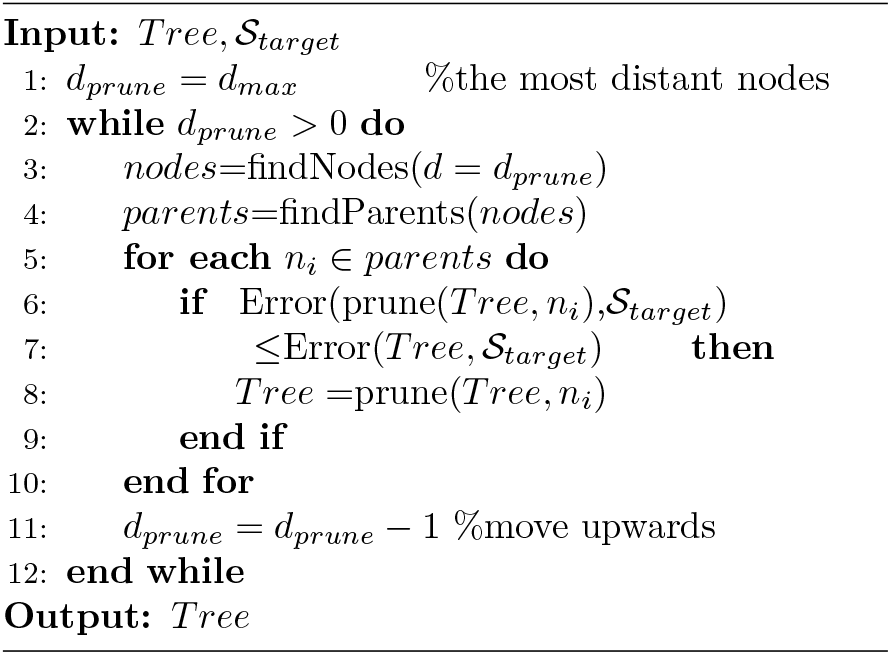

##### Algorithm 2 RF Self-Calibration

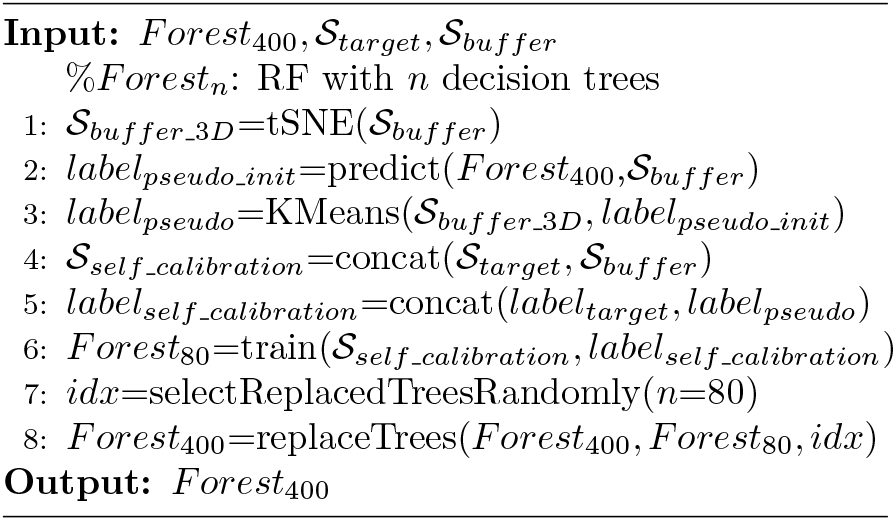

After decision tree pruning, we trained 200 additional trees from scratch, using the calibration data from only the new target user. These additional trees were appended to the pre-trained and pruned RF. The fine-tuned RF model comprises 400 decision trees in total.

### 2.4. Self-Calibration

The whole framework of self-calibration is presented in Figure 2. We used a data buffer to store the incoming testing data. The maximal size of the applied data buffer is 1500 windowed EMG feature vectors (~500 KB if using float32 format). Once a testing block completed, the data buffer was updated, and then used in the self-calibration process. When the buffer reached the maximal size, we deleted the oldest sample within the gesture category (pseudo-labels) with the most samples. Accordingly, samples in the data buffer tend to be balanced across different hand gestures. To self-calibrate a model, manifold learning and clustering were employed to assign pseudo-labels on buffered samples, which were then used to update the model parameters.

**Figure 2:**
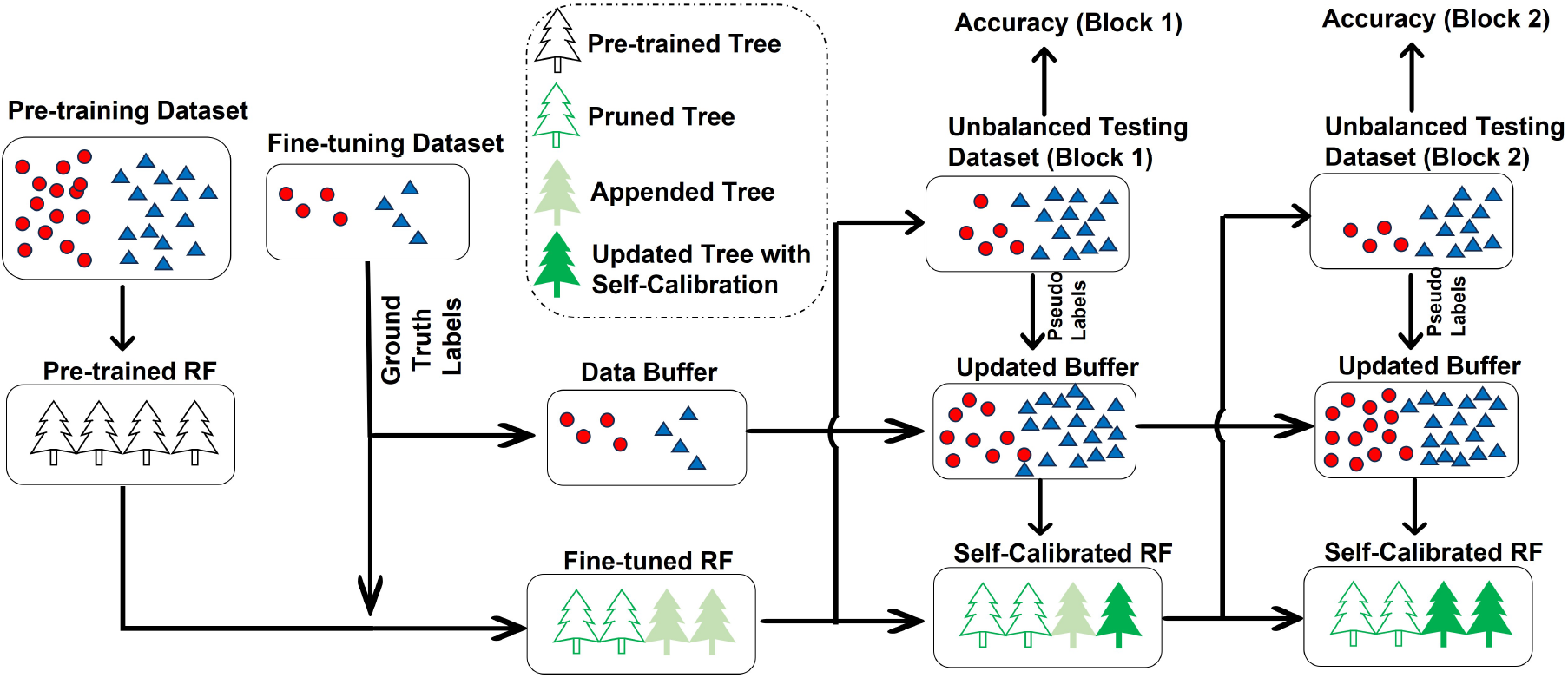
The framework of self-calibrating RF common model.

#### 2.4.1. Manifold Learning and Clustering

To assign more reliable and accurate pseudolabels on EMG data, dimensionality reduction was first performed. To better embed the complex distribution structure of EMG features, the dimensionality reduction enabling manifold learning was applied. Specifically, we employed t-Distributed Stochastic Neighbor Embedding (t-SNE) [33] to map the original high-dimensional (8 EMG channels *×* 10 types of features) EMG feature space into a 3-dimensional subspace. We applied t-SNE due to its capability to simplify the non-linear data distribution via manifold learning and meanwhile preserve the local structure of the original distribution. After dimensionality reduction via t-SNE, K-Means clustering was applied in the 3-dimensional t-SNE subspace. The initialisation of K-Means labels before iterations were determined as the classification outcomes given directly by the current RF model. After pseudo-label assignment, the number of samples corresponding to different classes (according to the pseudo-labels) might be different. The number of latest samples in the data buffer used in the following model updating step was determined by the minimal number of samples in each class.

#### 2.4.2. Updating Model Parameters

The calibration data (one repetition per gesture) with ground-truth labels, denoted as *S*_*target*_, together with data and pseudo-labels in the data buffer, denoted as *𝒮* _*buffer*_, were integrated together to train 80 decision trees from scratch and replace the original ones. Considering the pruned decision trees were pre-trained on data from other users which could provide different information with the data from the target user, we kept these pruned decision trees and replaced only the appended decision trees (randomly selected). To prevent the model from forgetting previous knowledge, each time we only replaced 80 appended decision trees. The pseudo-codes of RF self-calibration are presented in Algorithm 2.

### 2.5. Baseline Machine Learning Models

Two baseline machine learning models were implemented for comparison. First, a user-specific LDA model was trained from scratch using data from only the new target user. LDA was selected here because it is generally considered by the myoelectric community as a gold standard for the state-of-the-art models [34, 35, 36]. Second, a user-specific RF model (also with 400 decision trees) was trained from scratch using data from only the new target user. By comparing the performance of the pre-trained and fine-tuned RF model with the two baseline models, we aim to demonstrate the superiority of our model in achieving a higher initial accuracy without self-calibration. By further comparing model performances with and without self-calibration, we aim to demonstrate the effectiveness of the self-calibrating RF model.

### 2.6. Validation Methods

We performed both offline and real-time validations. The offline validation was performed using the two datasets collected in experiments 1 and 2. Data from all 20 participants in dataset 1 were allocated into the pre-training dataset. A leave-one-participant-out cross-validation was then applied on data in dataset 2 (18 participants). Each of the 18 participants was viewed as the testing participant in turn, with data from all the other 17 participants allocated as an additional pre-training dataset. Accordingly, data from 20+17=37 participants were used to pre-train a model for each testing participant. Data recorded from the target testing participant in the calibration section (1 repetition per gesture with 1s holding duration) were used to further fine-tune the pre-trained RF via decision tree pruning and appending. Once pre-trained and fine-tuned, the model was duplicated into two copies, one with and the other without self-calibration. For the self-calibrating model, self-calibration was performed to update model parameters after each testing block, i.e. when 20% of samples within the data buffer were updated. The baseline user-specific RF and LDA models were also trained (from scratch) using the same data collected from the calibration section. All these models will be compared in our offline analyses.

Considering the pre-trained and fine-tuned RF model outperformed user-specific RF and LDA models in offline analyses (see Section 3.1), in our real-time experiment, we only ran two models, i.e. the fine-tuned RF with and without self-calibration, in an open-loop mode (experiments on day 1 and the part 1 on day 2). In experiments of the part 2 on day 2, participants were divided into two groups. For participants in Group B, visual feedback corresponding to the self-calibrating model was presented. All hyper-parameters were justified in our offline analyses and directly used in the real-time experiment without any further hyper-parameter tuning.

### 2.7. Investigating the Effect of Visual Feedback

To explore the differences between Groups A and B (with and without visual feedback, respectively), we compared both the average classification accuracy and the EMG amplitude in our real-time experiments. To compare the effect of visual feedback on EMG amplitude, we first normalised the EMG amplitude (quantified by RMS) of all participants in part 1 on day 2 (before grouping, all in open-loop mode) to the same baseline level. Then the average amplitudes of participants with and without feedback in experiments of part 2 on day 2 were compared.

### 2.8. Investigating the Effect of Noises

We simulated the factor of noises by injecting Gaussian noises into the collected EMG data. In real-world applications using sparse electrodes (e.g. 8 electrodes in our work), one corrupted electrode due to strong noises would substantially degrade the performance of most models. The noises were injected to only testing data because the collection of pre-training and fine-tuning datasets can be supervised to avoid EMG signals in extremely low quality. For the testing data collected on day 1 and day 2, we randomly selected one corrupted electrode (not necessarily the same on two days). Signals in the corrupted channel were injected with additive white Gaussian noises with a signal-to-noise ratio (SNR) of 20 dB, 15 dB, 10 dB and 5 dB.

### 2.9. Ablation Analyses

The key step for self-calibration is assigning pseudo-labels to saved testing data. In our method, t-SNE-based manifold learning and clustering were applied to assign pseudo-labels. To show the necessity and contribution of each individual component, we performed an ablation experiment with each component removed from the processing pipeline. When removing t-SNE, clustering was directly applied to the original feature space. When removing clustering, pseudo-labels were directly given as the outcome of the model.

### 2.10. Statistical Analyses

To compare the effect of feedback on two groups of participants, the Kruskal–Wallis test was applied. To compare the performances of different models on the same group of participants, the Wilcoxon signed-rank test and the Friedman test were performed for the comparison with 2 and *≥* 3 models, respectively. For the Friedman test, the Nemenyi post-hoc test was further performed to evaluate the pair-wise difference. The significance level was set as *p <* 0.05.

## 3. Results

### 3.1. Results in Offline Analyses

Results in offline analyses are presented in Figure 3. According to Figure 3(a) and Figure 3(b), pruned and appended RF model contributed to a higher accuracy in comparison with standard participant-specific LDA and RF models on both days. Self-calibration further could improve the accuracy of pruned and appended RF. In addition, as presented in Figure 3(a), the self-calibrating RF model contributed to a slowly increasing accuracy on each day, demonstrating that the self-calibrating RF model could automatically adapt to the varying EMG characteristics. Results in the ablation experiment were presented in Figure 3(c). With either clustering or t-SNE removed, the self-calibrating model achieved a relatively lower accuracy, demonstrating the necessity of each component in the whole framework.

**Figure 3:**
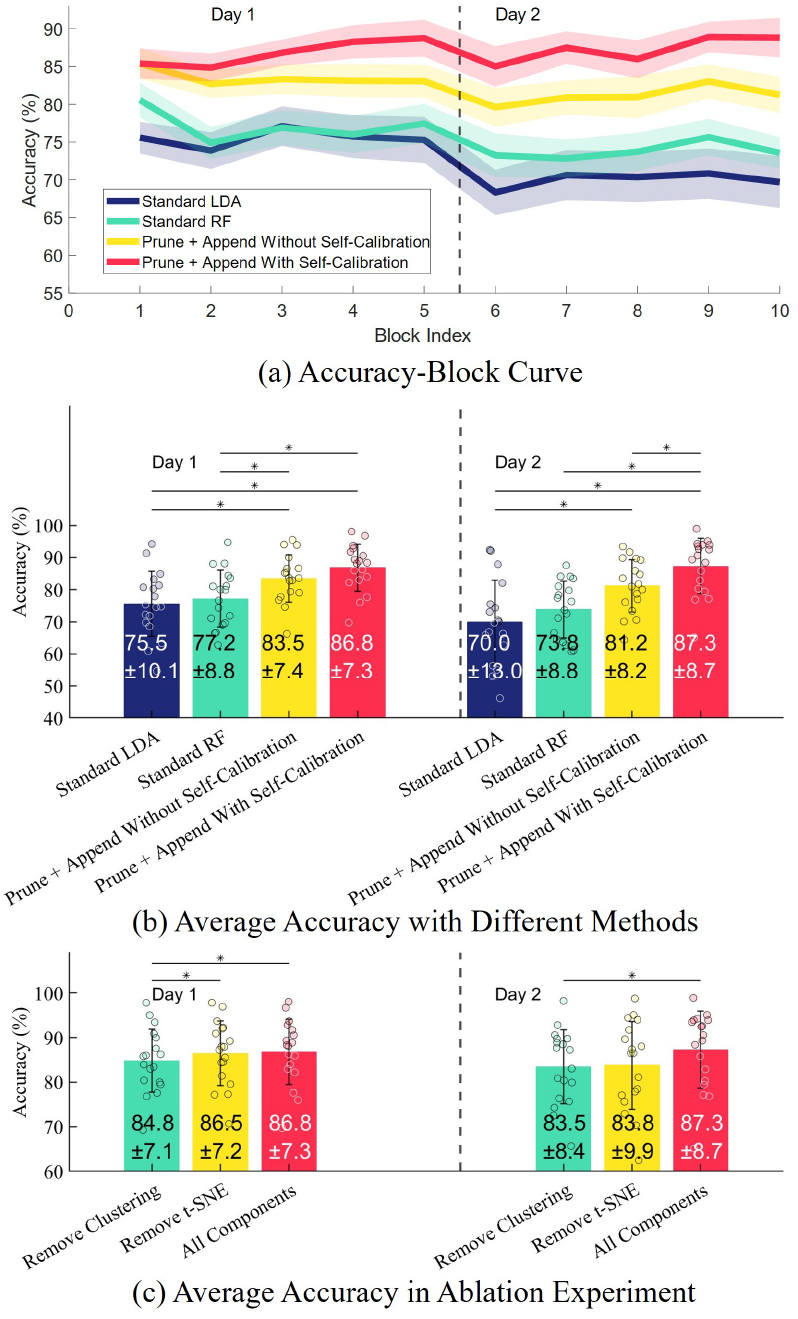
Results in offline analyses. (a) The offline classification accuracies of different models in different testing blocks (10 blocks on 2 days, with 5 blocks each day). For the figure clarity, standard error was shown as the shaded area. (b) The accuracy bar with standard deviation of each model. (c) The accuracy bar with standard deviation in the ablation experiment when each module is removed from the processing pipeline. In (b) and (c), the symbol “*” denotes a significant difference given by the Nemenyi post-hoc test of a Friedman test.

Figure 4 presents the accuracy variation with decreasing SNR. RF models showed a higher robustness against increasing levels of noise, compared with LDA. The pruned and appended RF model with self-calibration achieved the highest accuracy in all cases, although the improvement tended to decrease with stronger noises.

**Figure 4:**
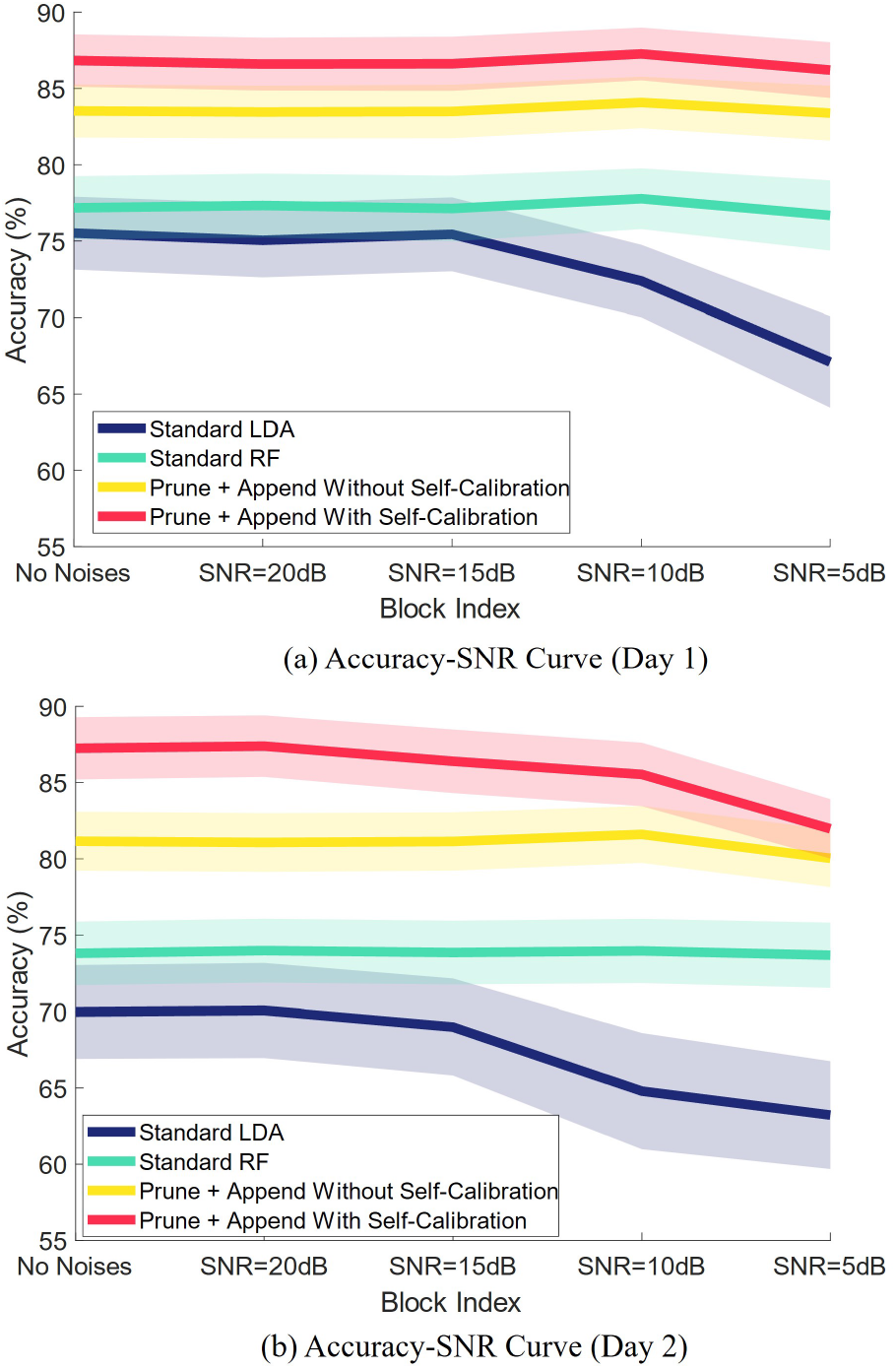
The effect of noises on classification accuracies. Two sub-figures (a) and (b) present results on day 1 and day 2, respectively. Standard error was presented as the shaded area.

### 3.2. Results in Real Time Experiments

Results in our real-time experiment are presented in Figure 5. According to Figure 5(a) and Figure 5(b), pruned and appended RF with self-calibration significantly outperformed the model without self-calibration on both day 1 and part 1 of day 2. As shown in Figure 3(a), the accuracy of the model without self-calibration progressively dropped from 84.3% (the first block on day 1) to 78.6% (the last block in part 1 on day 2). By contrast, the accuracy of the self-calibrating model progressively increased on day 1 (from 84.3% to 86.2%), then dropped at the beginning of day 2 (from 86.2% to 82.8%), and finally slowly increased by the end of part 1 on day 2 (from 82.8% to 86.5%).

**Figure 5:**
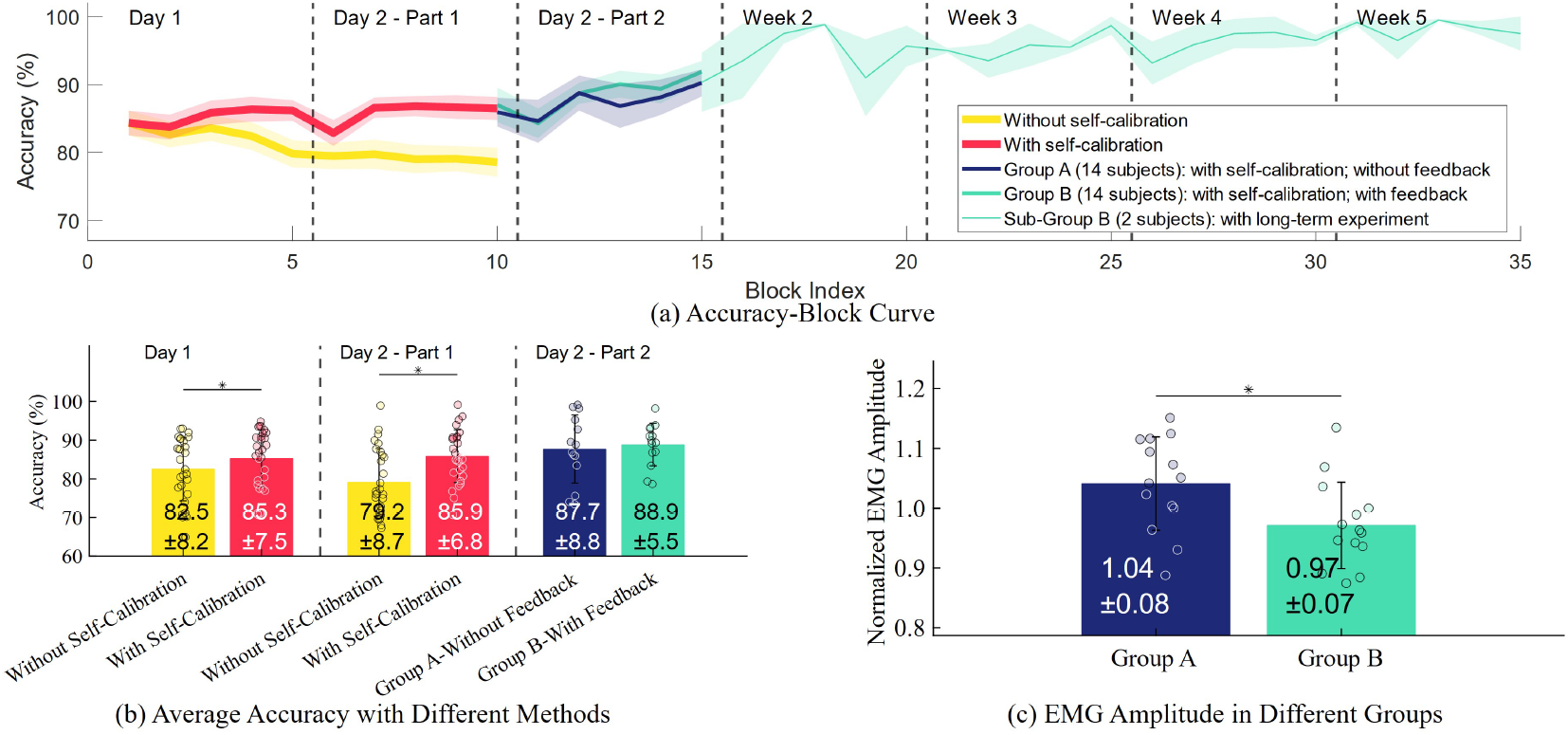
Results in real-time experiments. (a) The real time classification accuracies of different models in different testing blocks. Blocks 1–5: the open-loop experiment on day 1. Blocks 6–10: the open-loop experiment of part 1 on day 2. Blocks 11–15: the mixed open-loop and closed-loop experiments of part 2 on day 2, with participants separated in two groups. Blocks 16–35: the long-term closed-loop experiment lasting 4 more weeks for two participants in group B. For the figure clarity, standard error was shown as the shaded area. (b) The accuracy bar with standard deviation of each model. (c) The EMG amplitude bar (quantified by RMS) with standard deviation of each group (group A: open-loop experiment without feedback; group B: closed-loop experiment with feedback). In (b) and (c), the symbol “*” denotes a significant difference, given by the Wilcoxon signed-rank test and the Kruskal–Wallis test in (b) and (c), respectively.

In addition to the accuracy of the applied model in each block, we also show the accuracy of pseudo labels of the testing samples in the data buffer used for model self-calibration after each block (we show the results in the first 10 blocks in the open-loop experiment). As presented in Figure 6, the accuracy of pseudo labels show a similar trend as the accuracy of the calibrated model, with the accuracy increasing on day 1, dropping at the beginning of day 2, then gradually increasing on day 2 again. The overall average accuracy of pseudo labels is 84.4%.

**Figure 6:**
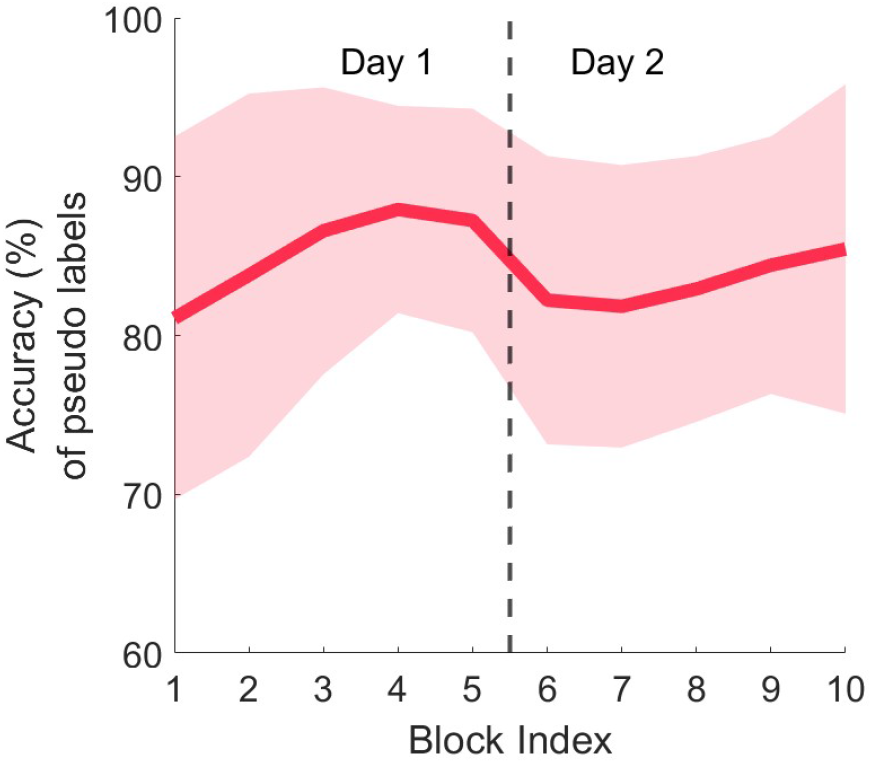
Accuracy of pseudo labels of the testing samples in the data buffer used for model self-calibration after each testing block. Standard deviation was presented as the shaded area.

We further compare the results between open and closed-loop experiments. According to Figure 5(a), in part 2 on day 2, the average accuracy of Group A (87.7%, without visual feedback) and Group B (88.9%, with visual feedback) did not show a significant difference. However, the EMG amplitude of participants in Group B with visual feedback was significantly lower than that of Group A without visual feedback, demonstrating that participants tended to learn the most effort-saving way to perform hand gestures with the help of feedback.

From Group B, we randomly selected two participants to take our 5-week long-term experiment. According to the curve in Figure 5(a), the accuracy can be maintained at a very high level. From week 1 (part 2 on day 2) to week 5, the average accuracy was 91.9%, 95.3%, 95.7%, 96.1% and 97.9%, even gradually increasing in long-term use. In the last week, the extremely high accuracy of 97.9% demonstrated the exciting prospects of our model in real-world applications.

## 4. Discussion

In this work, we aim to develop a plug-and-play myoelectric control system. We first pre-train a common RF model on data from other participants and then finetune the model using only one-repetition per gesture from the target testing participant. However, the large variability of EMG characteristics which may change over time degrade the performance of the model, due to extrinsic factors such as noises, electrode shift etc. Additionally, even without the extrinsic factors, the human motor system is inherently complex. Even when a myoelectric user repeats the same hand gesture, muscle activity can differ vastly, which is an inherent factor leading to degraded model performance. In addition, myoelectric users may gradually forget the details (e.g. the force level, the finger angles or the shape of their hand) of each hand gesture they have performed in the calibration session. Overall, using data from only one-repetition per gesture is quite challenging to capture the general EMG characteristics of a new user. As presented in Figure 5, the accuracy of the model without self-calibration gradually decreases even on the same day (blocks 1–5 on day 1), demonstrating the progressively increased level of distribution shift during the experiment. By contrast, in Figure 5, the accuracy of the self-calibrating model shows an increasing trend on both day 1 and day 2. The self-calibrating model could adapt to the gradually shifted data distribution, and even learn a more generalised data distribution from the shifted data to further improve the model’s generalisability.

In our experiment, the self-calibration process was performed after each testing block so that 20% samples in the data buffer were updated compared with the last calibration. The self-calibration process took less than one 1 minute on a PC with CPU (11th Gen Interl(R) Core(TM) i5-1145G7 @ 2.60GHz), without any parallel computing. With the inherent parallelisable advantage of RF model, the proposed self-calibration show promising prospects. In practical applications, the self-calibration can be performed once in a while when a certain number of samples in the data buffer are replaced by incoming new samples. The self-calibration can be performed in a background thread, without pausing the applications.

In our method, we assigned pseudo-labels by integrating t-SNE-based manifold learning and K-Means-based clustering. The high-dimensional EMG features are normally distributed on a curved manifold. This motivated us to apply manifold learning to simplify the high-dimensional data distribution. Manifold learning can flatten the curved high-dimensional manifold in a 3-dimensional subspace. The motivation of clustering is to jointly take into account (1) the knowledge already learned by the model (initialising the clustering labels before iterations) and (2) the distribution structure of a batch of stored data in the buffer. If we directly set pseudo-labels as the prediction outcomes of the current model without t-SNE manifold learning and K-Means clustering, the model may likely fall into a loop of learning biased knowledge given by itself. Our ablation experiment also demonstrated the necessity of manifold learning and the clustering steps. With simulated noises injected into an EMG channel, the model with self-calibration achieved the highest accuracy in all cases. However, it is noteworthy that with extremely strong noises, the performance improvement due to self-calibration tends to be less obvious. Models with and without self-calibration tend to reach similar performances at higher noise levels. As discussed before, self-calibration can adapt the model to a slowly shifted data distribution within a short period of time. Although the data distribution may substantially shift in long-term applications, the dynamic and progressively shifted data distribution during the application process serves as a bridge to connect the original and the final EMG patterns, so that the difficulty of each calibration could be reduced, contributing to a largely improved performance. Accordingly, it is foreseeable that there are extremes of the self-calibration mechanism. When the data distribution suddenly and vastly shifted, the benefits of the self-calibration mechanism would be reduced. That is why with extremely strong noises, the performance improvement due to self-calibration is reduced.

The concepts of supervised model fine-tuning [37], semi-supervised/unsupervised model calibration via pseudo-labels [20, 21, 22], and incremental learning [38, 39] have been explored in myoelectric control applications. Most previous methods were implemented on a deep neural network, due to its flexibility in model architecture and capability to be progressively trained/calibrated via gradient descent on network parameters. However, a deep network is commonly used as a black-box while explainable models are expected in human-centered and biomedical applications. Furthermore, the inference process of a deep network requires a large amount of floating point operations, increasing the computational burdens on low-cost mobile computing devices (e.g. microcontroller).

Our work differs with previous related studies from the aspects of both model architecture and validation methods. First, from the aspect of model architecture, we performed both fine-tuning and self-calibration on a pre-trained RF model. Conventional non-neural network machine learning models are generally considered incapable of being progressively trained, and can only be trained from scratch. Previous studies [40, 41] applied boosting-based methods to improve the robustness of non-neural network models by integrating multiple small base models. However, models in these previous studies [40, 41] were still trained in a user-specific way and evaluated offline. In this work, we prove that RF models are pre-trainable and can be fine-tuned and self-calibrate itself progressively. Given that RF model is also explainable [42], robust to a small size of training data [43], robust to noises [42], and parallelisable, the proposed self-calibrating RF provides a promising alternative for the myoelectric control community. Second, from the aspect of validation methods, previous studies validated the effectiveness of the model calibration algorithms in open-loop settings [17] or closed-loop settings but with EMG-independent-feedback [24]. In practical closed-loop applications, with decoder-specific feedback on classification outcomes presented to users, EMG characteristics would also progressively change and be biased to the specific implemented EMG decoder due to user motor learning. The performance of a model adapting to dynamic user behaviors and the feasibility of user adapting to dynamic self-calibrating model have not been well studied before.

The effect of feedback has been demonstrated to largely impact the performance of myoelectric control [44, 45]. Visual feedback has been proven effective in facilitating motor learning of users so that users can learn useful myoelectric skills and adapt to a fixed model, contributing to improved performance compared with that in an open-loop mode [45]. In this work, we explore the mechanism of bidirectional user-model co-adaptation when users interact with a dynamic self-calibrating model (rather than a fixed model). We find that, compared with user adaptation, the model adaptation via self-calibration can make the greatest contribution from the aspect of improving accuracy. With self-calibration, the model performance was significantly improved even in an open-loop mode. Further proving visual feedback can only slightly improve the performance (see Group B in Figure 5). In addition to the aspect of accuracy, another interesting finding is that visual feedback can reduce the amplitude of users’ EMG. With visual feedback, users can learn the most efficient manner to perform hand gestures and save efforts in long-term applications, which is another benefit of feedback for myoelectric control.

## 5. Conclusion

In this work, we propose a self-calibrating RF model, which can be first pre-trained on data from a large number of users and then fine-tuned using only one-repetition (1s signal duration) per gesture from the target user. The fine-tuned RF model can further self-calibrate itself in long-term use. Both offline and realtime, open and closed-loop, intra-day and long-term experiments demonstrate the robustness of our model. We explore the bidirectional user-model co-adaptation and find that visual feedback can help users learn a more efficient and effort-saving way to perform hand gestures. Our work contributes to the use of a plug-and-play myoelectric control model in real-world applications.

## 5.1 Acknowledgments

This work was supported by a grant from Engineering and Physical Sciences Research Council (EPSRC), UK (Grant No. EP/R004242/2).

